# Multi-omic analysis along the gut-brain axis points to a functional architecture of autism

**DOI:** 10.1101/2022.02.25.482050

**Authors:** James T. Morton, Dong-Min Jin, Robert H. Mills, Yan Shao, Gibraan Rahman, Daniel McDonald, Kirsten Berding, Brittany D. Needham, María Fernanda Zurita, Maude David, Olga V. Averina, Alexey S. Kovtun, Antonio Noto, Michele Mussap, Mingbang Wang, Daniel N. Frank, Ellen Li, Wenhao Zhou, Vassilios Fanos, Valery N. Danilenko, Dennis P. Wall, Paúl Cárdenas, Manuel E. Baldeón, Ramnik J. Xavier, Sarkis K. Mazmanian, Rob Knight, Jack A. Gilbert, Sharon M. Donovan, Trevor D. Lawley, Bob Carpenter, Richard Bonneau, Gaspar Taroncher-Oldenburg

## Abstract

Autism is a highly heritable neurodevelopmental disorder characterized by heterogeneous cognitive, behavioral and communication impairments. Disruption of the gut-brain axis (GBA) has been implicated in autism, with dozens of cross-sectional microbiome and other omic studies revealing autism-specific profiles along the GBA albeit with little agreement in composition or magnitude. To explore the functional architecture of autism, we developed an age and sex-matched Bayesian differential ranking algorithm that identified autism-specific profiles across 10 cross-sectional microbiome datasets and 15 other omic datasets, including dietary patterns, metabolomics, cytokine profiles, and human brain expression profiles. The analysis uncovered a highly significant, functional architecture along the GBA that encapsulated the overall heterogeneity of autism phenotypes. This architecture was determined by autism-specific amino acid, carbohydrate and lipid metabolism profiles predominantly encoded by microbial species in the genera *Prevotella, Enterococcus, Bifidobacterium*, and *Desulfovibrio*, and was mirrored in brain-associated gene expression profiles and restrictive dietary patterns in individuals with autism. Pro-inflammatory cytokine profiling and virome association analysis further supported the existence of an autism-specific architecture associated with particular microbial genera. Re-analysis of a longitudinal intervention study in autism recapitulated the cross-sectional profiles, and showed a strong association between temporal changes in microbiome composition and autism symptoms. Further elucidation of the functional architecture of autism, including of the role the microbiome plays in it, will require deep, multi-omic longitudinal intervention studies on well-defined stratified cohorts to support causal and mechanistic inference.

## Introduction

Autism spectrum disorder (ASD) encompasses a broad range of neurodevelopmental conditions defined by heterogeneous cognitive, behavioral and communication impairments that manifest early in childhood [1]. To date, over a hundred genes have been identified as putatively associated with ASD, with some genotypes now having a standardized clinical diagnosis [2]. However, most of the genetic variants are still associated with heterogeneous phenotypes, making it difficult to identify molecular mechanisms that might be responsible for particular impairments [3]. Some studies have also looked at the presence of abnormalities in different brain regions in children with ASD [4, 5]. However, whether such neuroanatomical features could mechanistically determine autism, and whether environmental factors could induce analogous ASD-like symptoms, remains unresolved [1].

In addition to risk factors, one comorbidity that has been linked to ASD with high confidence is the occurrence of gastrointestinal (GI) symptoms, such as constipation, diarrhea, or abdominal bloating, but causal insights remain elusive [6, 7, 8]. Mechanistically, much research has been focused on the interplay between the GI system and processes controlled by the neuroendocrine, neuroimmune, and autonomous nervous systems, all of which converge around the GI tract and together modulate the gut-brain axis (GBA) [9, 10, 11].

The GBA facilitates bidirectional communication between the gut and the brain, contributing to brain homeostasis and helping regulate cognitive and emotional functions [9, 12]. Over the past decade, research on the factors modulating the GBA has revealed the central role played by the gut microbiome—the trillions of microbes that colonize the gut—in regulating neuroimmune networks, modifying neural networks, and directly communicating with the brain [13]. Dysregulation of the gut microbiome and the ensuing disruption of the GBA are thought to contribute to the pathogenesis of neurodevelopmental disorders including autism, but the underlying mechanisms and the extent to which the microbiome explains these dynamics is still unknown [14, 15, 16, 17].

Several dozen autism gut metagenomics studies have revealed many, albeit inconsistent, variations in microbial diversity in individuals with ASD compared with neurotypical individuals [18, 19, 17]. Similarly, metagenome-based functional reconstructions and metabolic analyses have also shown strong, albeit inconclusive differences between ASD and neurotypical individuals [20, 21, 22]. Comparative analyses at other omic levels have further shown little agreement across studies [23] raising the question of whether the results obtained so far reflect intrinsic biological differences among cohorts, insufficient statistical power, or experimental biases that preclude meaningful comparisons [24].

A wide range of factors could explain the disagreement across studies, including confounding variation due to batch effects, the application of inappropriate statistical methodologies, and the vast phenotypic and genotypic heterogeneity of ASD. Batch effects can be caused by many factors including misspecified experimental designs, technical variability, geographical location, and demographic composition, and several algorithms have been proposed to correct for them, but a lack of standardized statistical methods further complicates interpretation [25, 26, 27, 28]. Microbiome datasets, like other omic datasets, are compositional, and failure to account for the compositional nature of sequencing counts can lead to high false positive and false negative rates when identifying differentially abundant microbes [29, 30, 31]. Microbiome analysis in ASD is further confounded by the phenotypic and genotypic heterogeneity of the disorder, which is known to be critical for stratifying ASD subtypes and constructing reliable diagnostics but is typically not measured or controlled for [32, 33, 1].

Understanding functional architecture—the network of interactions among different omic levels that determines individual phenotypes—of complex neurodevelopmental disorders such as autism, requires an accurate and comprehensive characterization of the different omic levels contributing to it [34]. Traditionally focused on the human genomic, metabolic, and cellular components of phenotype determination, mounting evidence of the role the GBA plays in phenotype determination through bidirectional modulatory mechanisms raises the need for considering the metagenomic and metabolic contributions of the microbiome as potential key components of the functional architecture of autism [35, 36].

To identify autism-specific omic profiles while reducing cohort-specific confounding factors, we have devised a Bayesian differential ranking algorithm to estimate a distribution of microbial differentials, or relative log-fold changes, [31] across multiple potential ASD subtypes implicit in 25 omic datasets (Table 1). A key feature of this approach was to match individual study participants by sex and age within each study to adjust for confounders in childhood development and cohort-specific batch effects. The preponderance of autism among males is well documented and several potentially sex-dependent mechanisms to explain this phenonmenon have been proposed [37]. Furthermore, the development of the microbiome during childhood is a hallmark of microbiome dynamics in the human gut [38, 39, 40, 41, 42]. Our analysis provides insights into the complexity of the interplay among multiple omic levels in ASD, highlights the inherent limitations of cross-sectional studies for understanding the functional architecture of autism, and provides a framework for further studies aimed at better defining the causal relationship between the microbiome and other omic levels and ASD.

**Table 1:** ASD omic datasets included in this study All sequencing datasets were retrieved from the SRA. [124].

| Organism | Body type | Data type | Num Studies | Num Subject pairs | References |
| --- | --- | --- | --- | --- | --- |
| Human | Postmortem brain tissue | RNAseq | 4 | 49 | (Velmeshev et al. 2019; Wright et al. 2017; Herrero et al. 2020) SRP072713 |
| Human | Serum | Immune markers | 1 | 22 | (Zurita et al. 2020) |
| Human | Serum | Metabolome | 2 | 50 | (Needham et al. 2021; Kuwabara et al. 2013) |
| Human | Urine | Metabolome | 1 | 26 | (Noto et al. 2014) |
| Human | NA | Dietary survey | 1 | 26 | (Berding and Donovan 2019) |
| Human | NA | Behavioral survey | 1 | 28 | (Kang et al. 2019) |
| Microbial | Fecal | Metabolome | 2 | 43 | (Needham et al. 2021; Kang et al. 2018) |
| Microbial | Fecal | 16S amplicon | 10 | 346 | (Berding and Donovan 2019; Zurita et al. 2020; Dan et al. 2020; J. Zhu et al. 2021; Fouquier et al. 2021; Zou et al. 2020; Kang et al. 2019, 2017; Son et al. 2015; David et al. 2021) SRP299486 |
| Microbial | Fecal | Shotgun metagenomics | 3 | 83 | (Averina et al. 2020; M. Wang et al. 2019; Dan et al. 2020) |

## Results

The structure of our analysis consisted of a multi-cohort and multi-omic meta-analysis framework that allowed us to combine independent and dependent omic data sets in one integrated analysis [43, 44]. To minimize issues of compositionality and sequencing depth [45], we modeled overdispersion using a negative binomial distribution for modeling sequencing count data [46] (Box 1 “TACKLING METAGENOMIC UNKNOWNS”). Our differential ranking approach incorporated a case-control matching component consisting of individually pairing ASD children with age- and sex-matched neurotypical control children within each study cohort, allowing us to adjust for confounding variation and batch effects (Supplemental methods). Finally, we cross-referenced the 16S–based microbial differential ranking analysis from eight age-sex matched cohorts against 15 other omic datasets to contextualize the potential functional roles these microbes could play in autism (Figure 1).

**Figure 1:**
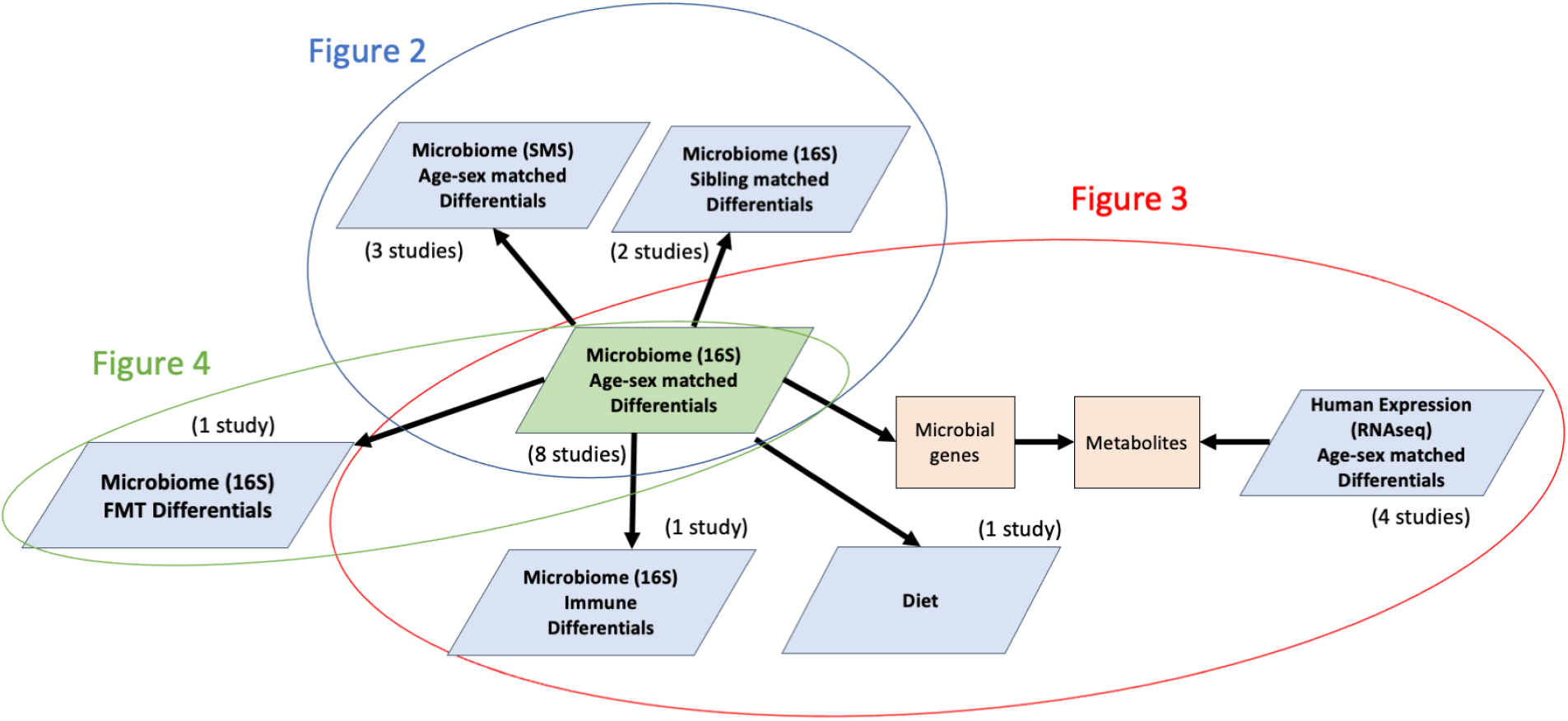
The structure of our meta-analysis across multiple omics levels. For Figure 2, microbial differential abundance on 16S data from age-sex matched cohorts was cross-referenced against sibling matched cohorts and SMS data from other age-sex matched cohorts. For Figure 3, these same 16S microbial differential abundances were cross-referenced against cytokine profiles, dietary surveys and pathways from RNAseq. For Figure 4, the differentially abundant microbes from the age-sex matched analysis was cross-referenced against the Kang et al FMT trial.

### Age- and sex-matching increases informational content of cross-sectional ASD datasets

We compared the age- and sex-matched differential ranking analysis to the standard group-averaged differential ranking analysis across eight out of the ten 16S studies [47, 48, 49, 21, 50, 51, 52, 53]. Age- and sex-matched differential analysis outperformed standard group averaging with respect to *R*^2^, and its overall performance strictly improved as more studies were added (Figure S1). This performance boost reflected a reduction in model uncertainty with larger cohorts that was indicative of overlapping differentially abundant taxa across studies and of reduced confounding variation.

### Global differential ranking analysis reveals a distribution of significant ASD-microbiome associations

A global, age- and sex-matched differential ranking analysis of the eight 16S datasets selected for this study revealed a clear partitioning of microbial differences with respect to ASD and cohort membership (Figure 2a, Figure S2). The distribution of the overall case-control differences showed a strong ASD-specific signal driven by 142 microbes more commonly found in ASD children and 32 microbes more commonly found in their control counterparts (Table S1). The variability observed is most likely due to confounding factors such as cohort demographics and geographic location, with the eight cohorts originating from Asia, Europe, South America, and North America. Analogous global differential ranking trends could be observed for the virome, shotgun metagenomics sequencing (SMS), and RNA-seq datasets (Figure S3). To determine whether these highly significant microbiome signals (pvalue*<*0.0025) could be used to distinguish ASD subjects from their age- and sex-matched control counterparts, we trained random forest classifiers on train/validation/test splits on data derived from 16S—targeted sequencing of the microbial 16S ribosomal RNA gene—and SMS—whole genome sequencing of microbial communities. Despite the strong microbiome effect size, we faced difficulties fitting generalizable classifiers. Our best classifiers had an average cross-validation accuracy of about 75% (Figure 2b), falling within the range of 52%–90% classification accuracy observed in previous studies [50, 49, 54]. We suspect that the vast heterogeneity across cohorts hampered classification performance. While cohort size did not impact predictiability (Figure 2c), some cohorts with skewed sex ratios or age ranges did exhibit lower classification performance (Figure. 2d-e). In the Zurita et al. cohort, sex-specific factors could confound classification (four girls and 56 boys) [48], and in the Kang et al. cohort, age-associated microbiome development factors could hamper classification accuracy (all children were 10 years or older). In addition, all subjects in the Kang et al. cohort had known GI symptoms [55], further compromising classifier performance because none of the other studies controlled for this variable. As a result, and analogous to the phenotypic and genotypic heterogeneity observed in ASD, the microbiome composition of ASD children also exhibits high heterogeneity, precluding the identification of a homogeneous universal ASD microbial profile and the construction of generalizable classifiers.

**Figure 2:**
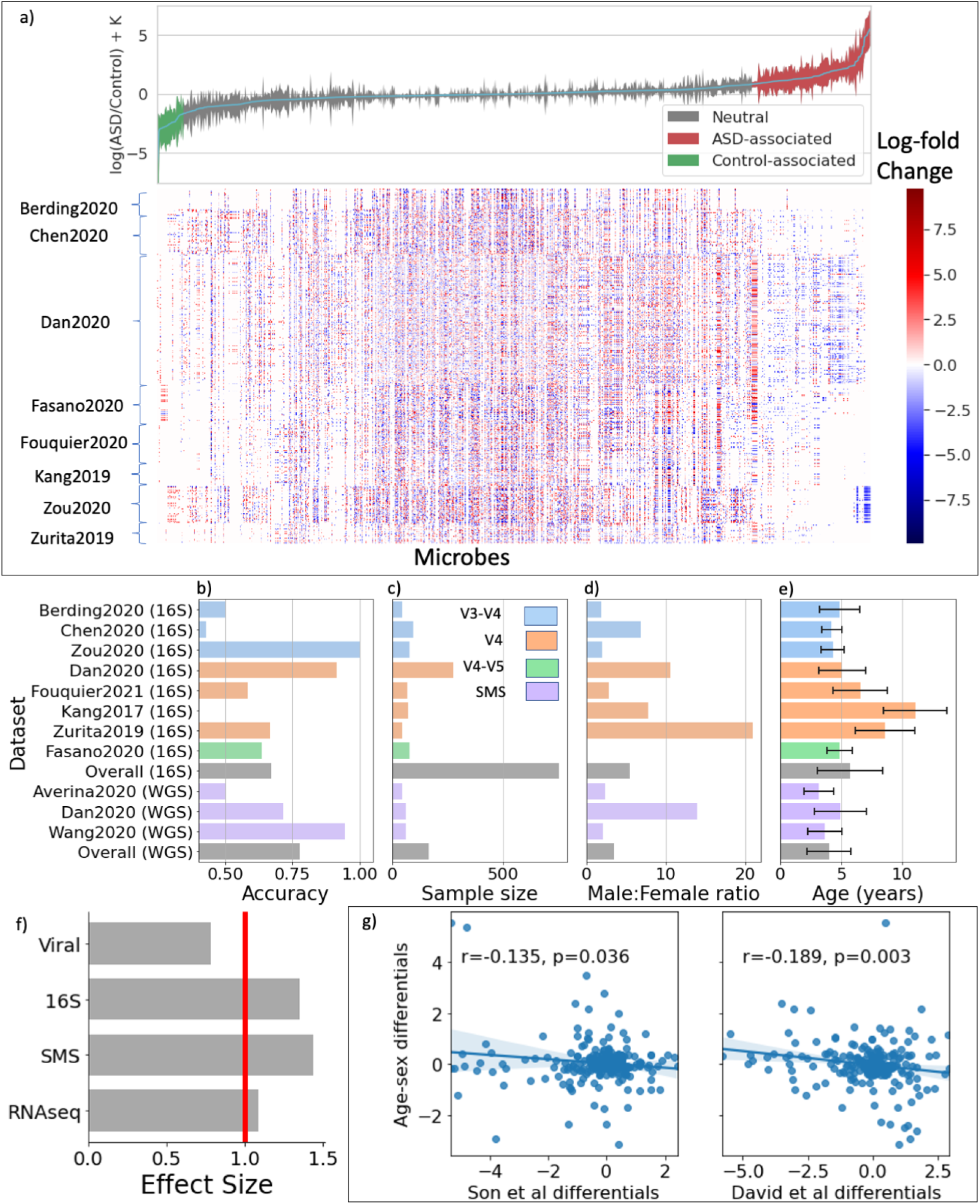
Differential ranking analysis across omics levels. (a) Global microbial 16S log-fold changes between age- and sex-matched ASD and control individuals. Error bars represent the 95% credible intervals. Heatmap showing all CLR transformed microbial differentials for each age- and sex-matched ASD-control pair across all cohorts. (b-e) Held-out random forests ASD classification accuracy, sample size, male:female ratio and age distributions across all 16S and shotgun metagenomics datasets analyzed in this study. V3-V4, V4, and V4-V5 refer to the variable region of the bacterial ribosomal RNA analyzed; SMS refers to shotgun metagenomic sequencing. (f) Effect sizes of different omics levels: viral, 16S, SMS, and RNAseq. (g) Comparisons of sibling-matched differentials from two different studies to the global age- and sex-matched differentials

### Children with ASD exhibit significant individual differences at several omic levels

Differential ranking analysis of three core omic levels—microbiome (16S and SMS) and human transcriptome (RNAseq)—revealed strong and highly significant differences between ASD subjects and their neurotypical counterparts (p-value *<* 0.0025) (Figure 2f; Table S2; Table S3). Two additional omic levels—the metabolome and the virome— didn’t show significance signals (Figure S4, Table S4). Amongst the models that yielded a statistically significant signal, the 16S and SMS datasets had a larger effect size than the RNAseq datasets (Figure 2f). While each omic dataset by itself showed strong associations with ASD, a side-by-side comparison of the 16S and SMS datasets—two datasets that should show high equivalence—revealed a significant lack of overlap between them, highlighting the outstanding challenge of batch effects in microbiome studies (Figure S5). The reasons for this discrepancy could be many, but most likely center around sample size—our study looked at eight 16S datasets versus only three SMS datasets containing 754 individuals and 166 individuals, respectively. Another major challenge when estimating species profiles with metagenomics reference libraries is assigning a species identification to a read—there are many reads that do not uniquely map to individual species—and as a result, these multi-mapped reads can give rise to numerous false positive taxa [56, 57, 58, 59]. Based on this, we decided to focus primarily on the 16S datasets to define a global differential ranking profile.

### Sibling-matching and unrelated sex- and age-matching show significant discrepancies

To determine whether sex- and age-matched differentials could be universally predictive, we compared the 16S differentials obtained from the age- and sex-matched cohorts with two sibling-matched cohorts [60] [61]. Interestingly, we observed a significant negative correlation between the differentials extracted from the age- and sex-matched cohort and the two sibling-matched cohorts, suggesting that ASD-specific microbes in the sibling-matched studies are enriched in the control group in the age- and sex-matched studies and vice versa (Fig. 2g). Permanova applied to the age-sex matched cohort revealed a strong age-confounder across cohorts (pvalue *<* 0.001), with little confounding variation due to sex (Table S5). In contrast, Permanova applied to these sibling-matched cohorts revealed that household is a major confounder in both cohorts (pvalue *<* 0.001), but ruled out age as a confounder and indicated that sex was a confounder only in the David et al. cohort (pvalue *<* 0.001). The observed discrepancies point to different sets of confounders possibly affecting the analysis: in the case of the age- and sex-matched studies, family confounders aren’t typically accounted for, while sibling-matched studies don’t typically adjust for age confounders. In addition, and while cohorts such as the one studied in Maude et al. specifically control for the possibility, siblings often exhibit a higher risk of developing ASD compared to the general population [62].

### Host cytokine concentrations are correlated with microbial abundances

Immune dysregulation, ranging from circulating ‘anti-brain’ antibodies and perturbed cytokine profiles to simply having a family history of immune disorders, has been repeatedly associated with ASD [63, 64]. Recently, for example, Zurita et al. showed that concentrations of the inflammatory cytokine transforming growth factor beta (TGF-*β*) are significantly elevated in ASD children. We reanalyzed this dataset, after age- and sex-matching, and observed that microbial differentials associated with TGF-*β* and IL-6 concentrations were positively correlated with the global microbial log-fold changes between ASD and control pairs (IL-6 : r=−0.435, p=0; TGF-*β* : r=0.291, p=0) (Table S6). To validate the integrity of these microbial profiles with respect to the cytokine changes, we calculated the log-ratios of these microbial abundances and showed them to, in turn, be highly correlated with TGF-*β* and IL-6 concentrations (IL-6 : r=0.50, p=0.0007; TGF-*β* : r=0.45, p=0.002) (Figure 3 a-d).

**Figure 3:**
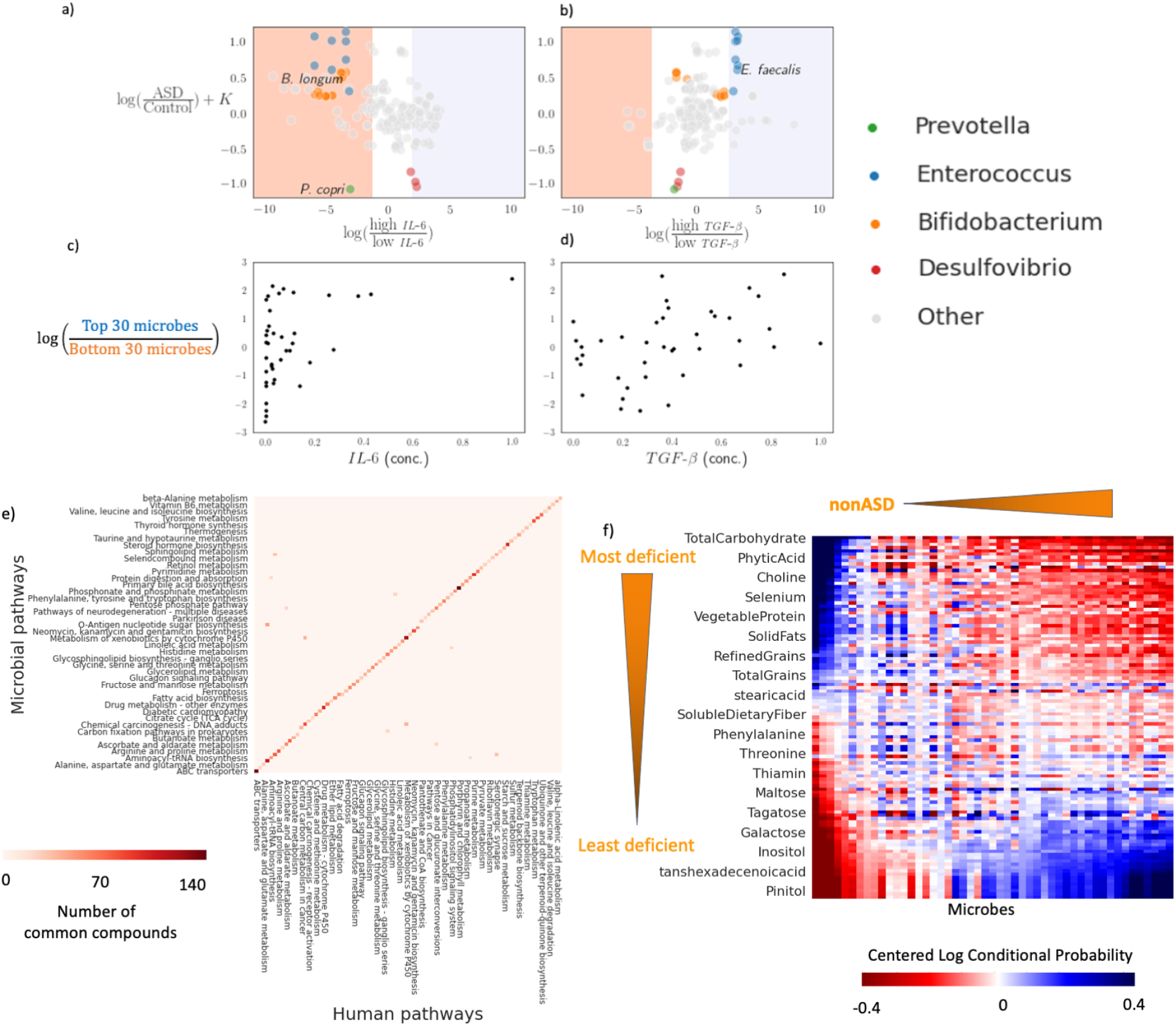
Characterizing the associations between differentially abundant microbes in ASD and cytokines, gene expression in the brain, and dietary patterns. (a-b) Comparison of microbial differentials obtained from age- and sex-matching and cytokine analysis. LFC denotes log-fold change of microbial abundances with respect to a specific cytokine. (c-d) Microbial log-ratios constructed from 30 top and bottom most differentially abundant microbes corresponding to each cytokine. (e) Heatmap showing the overlap of molecules between ASD-enriched pathways in the microbiome and in the brain. (f) Co-occurrence analysis between diet and microbes in ASD.

Four clusters of microbial genera—*Prevotella, Enterococcus, Bifidobacteria*, and *Desulfovibrio*—were predominantly associated with the cytokine differentials. Partial mechanistic insights on some of these cytokine-microbe associations have been previously published. Both *B. longum* and *E. faecalis* have shown antiinflammatory activities: *B. longum* downregulates IL-6 in fetal human enterocytes in vitro [65] and *E. faecalis* has been observed to upregulate TGF-*β* in human intestinal cells [66]. *P. copri* associations with different cytokines have also been observed in multiple disease contexts [67]. Similarly, *Bifidobacteria* and *Prevotella* both co-occurred with phages enriched in ASD or in neurotypical children (Figure S6, Table S7), but while microbes have previously been reported to mediate viral infections [68, 69], the mechanistic underpinnings of these interactions with the host’s immunity remain poorly understood [70, 71, 72].

### The microbiome metabolic capacity is reflective of the human brain-associated metabolic capacity in ASD

To determine potential crosstalk between the human brain and the microbiome physiology, we compared the metabolic capacities encoded by the microbial metagenome—combining the individual metabolic capacities of thousands of different microbes—and the differentially expressed human genome in the brain, two omic levels representing entirely different biological contexts. We observed that over 100 human metabolic pathways differentially expressed in the brain tissues of ASD individuals had analogous microbial pathways differentially abundant in the microbiome of children with ASD, suggesting a potential coordination of metabolic pathways across omic levels in ASD (Fig. 3e). Pathways related to amino acid metabolism, carbohydrate metabolism and lipid metabolism were disproportionately represented among the overlapping genes (Table S8).

### The microbiome metabolic capacity reflects restrictive diet patterns in children with ASD

Autistic traits in early childhood have been shown to correlate with poor diet quality later in life, however, little is known about how diet quality is directly linked to autistic traits [73]. Here, we re-analyzed the paired microbiome and dietary survey data from Berding et al.. A microbiome-diet co-occurrence analysis revealed startlingly similar amino acid, carbohydrate and lipid metabolism association patterns to those observed in the microbiome-brain metabolic capacity analysis (Table S9). Interestingly, both microbes enriched in ASD and in control subjects co-occurred primarily with amino acid dietary compounds (Figure 3f, Figure S7). Autistic children were less likely to consume foods high in glutamic acid, serine, choline, phenylalanine, leucine, tyrosine, valine and histidine, all compounds involved in neurotransmitter biosynthesis. Even though the metabolomic analysis did not yield statistically significant signals after FDR correction, the metabolites that showed the strongest signal included glutamate and phenylalanine, consistent with the microbiome-diet analysis [74, 75, 76]. Disruptions in the biosynthesis of these neurotransmitter molecules have been implicated in a wide variety of psychiatric disorders, and a recent blood metabolomics study has shown the potential of using branched chain amino acids to define autism subtypes [33]. Due to the incompatibility between the molecular features across datasets, it was not possible to combine any of the metabolomics datasets to boost the statistical power, which remains a major limitation of metabolomics technologies at present (see Methods).

### Differential microbial rankings show disease-specific correlations

One major challenge in determining microbiome-disease associations is identifying correlations specific to a particular condition and not generally present across diseases [77, 78]. To determine how specific to ASD our global differential microbiome profile was, we cross-referenced it against differential ranking results obtained from an Inflammatory Bowel Disorder (IBD) dataset [79] and a Type 1 Diabetes (T1D) dataset [80]. IBD shares some comorbidities with ASD [81, 82], while no direct correlation between ASD and T1D has been reported to date. The analysis revealed a notable overlap between microbes enriched in ASD and IBD, and this overlap was stronger than both the overlap between IBD and T1D and between ASD and T1D (Figure S8). Whether this ASD-IBD overlap is suggestive of a common microbial profile or is confounded by the restrictive dietary nature of these two clinical conditions is currently unclear. Higher resolution and properly designed clinical studies will have to be performed to get to a mechanistic understanding of the potential microbiome connection between these two conditions.

### ASD microbiome profiles weaken after fecal matter transplant consistent with reported behavioral improvement

While the preceding cross-sectional analyses showed significant associations among several omic levels (virome, microbiome, immunome) or diet and ASD, insights into causality are still limited. By contrast, longitudinal intervention studies provide an opportunity to obtain stronger insights into causality. To test this, we re-analyzed data from a two-year, open label fecal matter transplant (FMT) study with 18 children with ASD [83]. In this study, the children were subjected to a two-week antibiotic treatment and a bowel cleanse followed by two days of high dose FMT treatment and eight weeks of daily maintenance FMT doses. Based on one of the most common evaluation scales for ASD, the Childhood Autism Rating Scale (CARS), significant improvements were achieved after the ten week course of treatment. Two months later the initial gains were largely maintained, and a two-year follow-up showed signs of further improvement in most of the patients. The results are consistent with a potential role of the microbiome in improving autism symptoms, but how the underlying changes in microbiome composition related to those seen in other studies remained unknown.

Here, we re-analyzed the original raw data in the context of the ASD profiles revealed by our cross-sectional differential ranking analysis (Table S10). All microbes associated with ASD in the 18 children prior to the FMT treatment had been identified as ASD-associated microbes in our age- and sex-matched cross-sectional analysis, recapitulating 74% of the cross-sectional profile. Immediately following FMT treatment, the abundances of the ASD-associated microbes decreased in all 8 children (Figure 4). The two-year followup analysis revealed that all the ASD-associated microbes, mostly *Enterococcaceae*, continued to be depleted. Consistent with the findings of Kang et al., we also observed *Desulfovibrio sp*., and *P. copri* increase over the two year period, while *Bifidobacteria sp*. could be found both among depleted and enriched species and other *Prevotella sp*. were depleted, pointing to a potentially wide functional diversity within these genera not noted in the original study.

**Figure 4:**
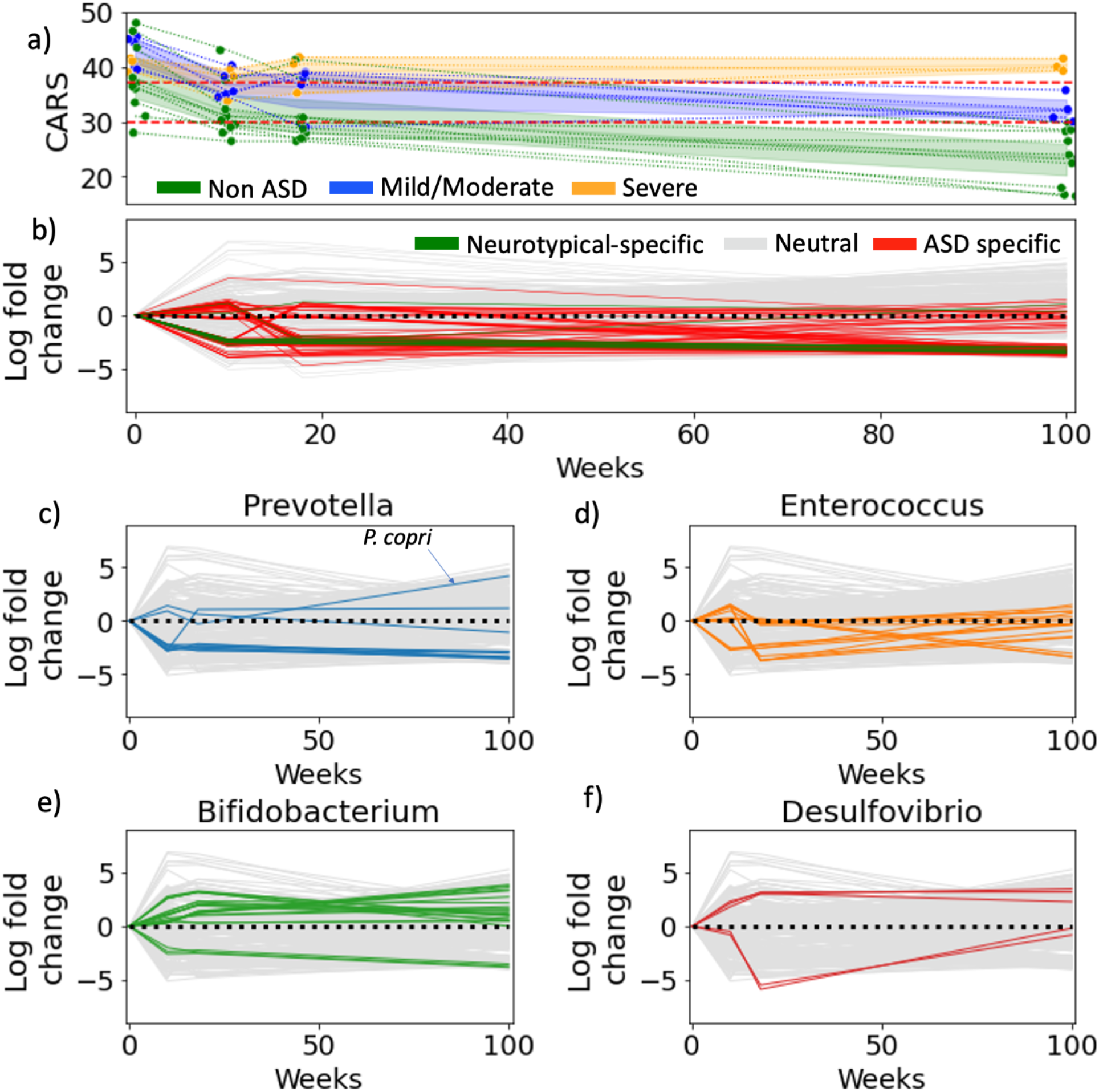
Fecal matter transplants have long-lasting effects on autism gut microbiomes. (a) The improvement of CARS for each ASD child over time. The children are split into 3 groups, non-ASD, mild/moderate and severe based on whether their CARS score fell below 30, between 30-37 or greater than 37 (b) Microbial log-fold changes over time: the time series was generated by calculating log-fold changes between time points for each microbe. ASD-specific microbes highlighted in red were determined in the cross-sectional study. (c-f) Microbial log-fold changes are re-colored with genera highlighted in cytokine comparisons.

## Discussion

The functional architecture of ASD, and in particular the potential role the microbiome plays in modulating the GBA in the context of autism, remains poorly understood due to disagreements among existing microbiome and other omic studies. Our Bayesian model highlighted a distribution of highly significant microbial differentials obtained from individual age- and sex-pairings between children with ASD and neurotypicals, and parallel analyses at the immunome, human transcriptome, and dietome levels revealed strong associations among omic levels. The virome and the direct metabolome signals, while present, were markedly weaker than the other omic signals. The inferred ASD-specific metabolic profiles from the microbiome and the human transcriptome, on the other hand, showed a high and significant degree of overlap in microbial and human pathways expressed in the gut and in the brain, respectively. The metabolic connection implied by this overlap, which included differentially enriched carbohydrate and amino acid metabolic pathways in ASD, is a remarkable observation given the fundamental difference between the gut and brain physiologies, which would a priori suggest a reduced overlap in metabolic capacities. The microbiome-diet co-occurrence analysis also highlighted a reduced intake of amino acids and carbohydrates linked to specific microbiome profiles in ASD children. These metabolic and dietary imbalances, particularly regarding glutamate levels, were further apparent, albeit weakly, in the serum, fecal and urine metabolomes we analyzed. This multiscale overlap we observed along the GBA points to the existence of a functional architecture of ASD driven by the metabolic potential at the genomic and metagenomic levels.

While the differential distributions we determined were highly significant, the global analysis did not provide reliable ASD classifiers or uncover universal microbial ‘smoking guns’ linked to autism. However, several microorganisms consistently detected across omic levels pointed to potentially interesting functional connections. For example, our analysis suggested that *B. longum* exhibited a down-regulation of IL-6, which has been observed across a number of in vitro and cohort studies[84, 85]. The diet co-occurrence analysis also showed a strong association between *P. copri* and carbohydrate depletion in ASD. The population dynamics of *P. copri* have been reported to be driven primarily by carbohydrates in the diet [67]. Multiple other microbes, including several *Bifidobacteria, Enterococcus* and *Desulfovibrio* species, stood out in the immune and viral analyses. In the FMT study, the relative proportions of several *Prevotella, Bifidobacteria, Desulfovibrio* and *Enterococcus* species also showed strong associations with ASD symptoms, further suggesting a causal role for these microorganisms in shaping autism symptoms.

Despite our inability to determine actual metabolomic profiles at this point (see Methods), our metabolite analysis based on microbiome- and brain-derived metabolite inferences as well as the diet-derived metabolite data reveals a picture of a unifying and distinct ASD functional architecture. With the brain, the immunome and diet as major effectors, the multi-factorial complexity of ASD is reduced to a multi-scale set of interactions centered around human and bacterial metabolism that in turn determines phenotypic, genomic and metagenomic attributes via multiple feedback loops (Figure 5). The association of specific genotypes with ASD has been clearly established [2]; the pivotal role of the immune system in mediating the communication between the gut microbiome and the human brain as well as other peripheral systems is also firmly established [86]; further, the central role of the microbiome in mediating diet-derived nutrient mobilization has been extensively documented [87]; and several hard-wired feedback loops among these effectors such as the hypothalamus-mediated regulation of appetite and diet, have also been described [10].

**Figure 5:**
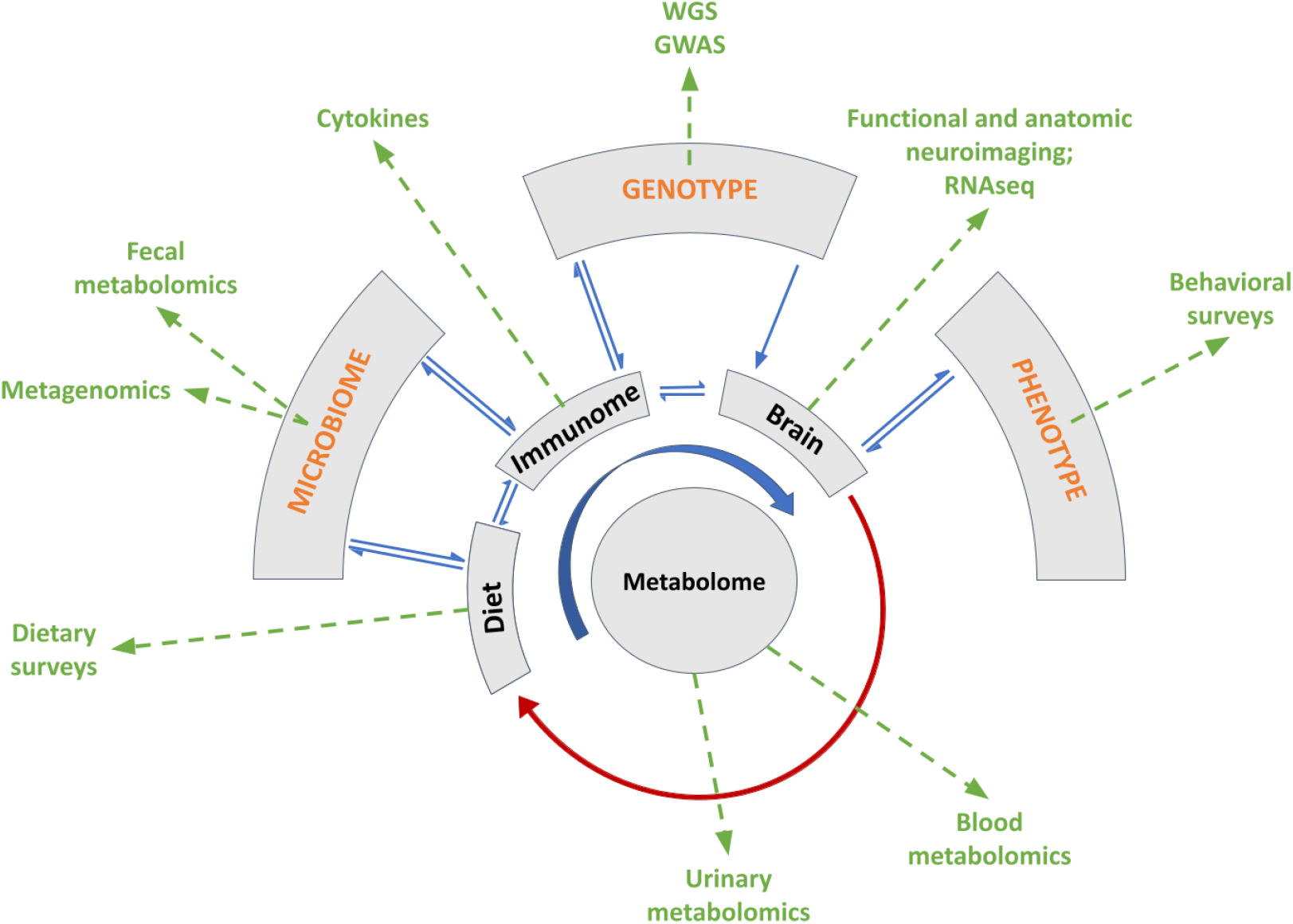
Proposed functional architecture of ASD Hypothesized causal graphs underlying relevant omic levels along the GBA in ASD, and experimental considerations for future studies. Blue arrows denote causal direction, red arrows indicate feedback loops and green arrows indicate measurable data types.

A major limitation of our meta-analysis is the lack of consistent behavioral, genotypic or electronic healthcare record data that would have allowed subtyping of the ASD subjects in light of environmental confounders. Furthermore, we cannot definitely recommend age- and sex-matching over sibling matching based on our analysis. Age remains a major confounding factor in early childhood microbiome development and controlling for this is key for understanding microbial fluctuations [88]. On the other hand, sibling-matching may help control environmental factors, but mostly rules out the ability to age-match subjects, thus potentially introducing the age confounder [89]. And while our approach revealed strong associations among the microbiome, other omic levels, and ASD, the vast heterogeneity in behavioral patterns and in genotypes is a major obstacle in constructing diagnostics and treatments for ASD symptoms [90, 32].

Our analysis has further exposed the limitations of cross-sectional cohort studies and the need for longitudinal intervention studies to further our understanding of the functional architecture of ASD. Building realistic causal models of autism needs to take into account the multi-factorial complexity underlying different ASD subtypes, which will require a concerted effort to simultaneously analyze several omic levels and at clinically relevant time scales. For instance, understanding the engraftment dynamics of FMT and its functional implications on the recipients’ gut microbiomes requires frequent initial sampling of the microbiome, immunome and metabolome, but tracing any behavioral changes over time requires less frequent sampling over periods of up to several years in combination with reliable behavioral, medical and dietary surveys [91, 92]. Collecting and integrating such multi-scale omic datasets presents unique logistical and analytical challenges.

Managing data acquisition and access will require coordinating multiple sites and potentially centralizing some aspects of sample processing. Recent initiatives such as The Environmental Determinants of Diabetes in the Young (TEDDY) study, an international long-term, multi-center initiative to link specific environmental triggers to particular Type 1 diabetes–associated genotypes, provide a blueprint for similar approaches in ASD [93]. A key component of such an initiative would be the establishment of standardized sampling and processing protocols that would minimize technical confounders, one of the top confounders at most omic levels. For instance, our analysis showed major batch effects when comparing 16S and SMS datasets across cohorts (r=−0.023, p=0.48, Figure S5b) as well as within a cohort (r=0.17, p=1e-5, Figure S5b). And while there are extensive efforts underway to calibrate microbiome datasets [94], other omic levels such as the metabolome [26] present even more fundamental technical issues that make it imperative to develop concerted strategies to be able to include them in an integrated analysis.

In addition to the considerable variations in statistical properties across datasets, interactions among omic levels are mostly underdetermined, making the construction of informative models a major challenge. Determining the necessary biologically relevant and unbiased assumptions is a non-trivial process and can inadvertently lead to model mis-specifications resulting in misleading conclusions. As pointed out recently for genetic, environmental and microbiome models in ASD [95, 96], addressing these issues will be critical to inferring causal mechanisms from population-scale studies. In addition, and given the vast heterogeneity of ASD, designing cohort studies that minimize confounding factor effects will be key to furthering our understanding of autism. For example, while our analysis could not identify ASD subtypes implicated in GI symptoms, we have determined stronger associations between gut microbes, host immunity, brain expression and dietary patterns than previously reported, highlighting the potential for boosting the statistical power and biological insight with comprehensive omic analyses. We conclude that multi-omic longitudinal intervention studies on appropriately stratified cohorts, in combination with comprehensive patient metadata, provide an optimal approach to advance our understanding of the etiology of autism to the next level.

## Methods

### Search strategy and inclusion criteria

We performed a systematic search for published and/or publicly deposited or not yet published and/or publicly available human microbiome, metabolome, immunome, transcriptome AND autism/ASD datasets in several NCBI databases (PubMed, SRA, and BioProject), UCSD’s MassIVE resource, the PsychENCODE consortium, the American Gut Project, and from individual research groups worldwide. About half of 70+ studies we identified were already deposited on public data repositories or were made directly available to us by the research groups.

Most studies consisted of heterogenous—no genotype or phenotype stratification—ASD and neurotypical age- and sex-matched cohorts and had one or two datasets (microbiome [16S, shotgun metagenomic sequencing (SMS)], metabolome [urine/serum/fecal], immunome [cytokines], transcriptome [RNAseq], dietary survey, behavioral survey) associated with them, with only a few studies having three or more omic datasets associated with them (Table 1). We adopted a multi-cohort and multi-omics meta-analysis framework that allowed us to combine independent and dependent omic data sets in one overall analysis[43, 44]. In total, we analyzed 597 ASD-control pairs. To reduce the batch effects and noise associated with primer choice in the 16S datasets, a major confounder in microbiome analyses, we restricted the 16S datasets to include only those targeting the variable region V4 of the bacterial ribosomal RNA, a region exhibiting higher heterogeneity and lower evolution rates than other variable regions[97, 98, 99]. Our analysis included 16S datasets obtained targeting the V4 region exclusively, the V3-V4 region, or the V4-V5 region.

The final metabolomic meta-analysis we present here consists of the combined analysis of only four independently preprocessed, normalized, and analyzed metabolomic datasets. Despite several more ASD-related datasets being available, the disparity in mass spectrometric technologies used to generate them, which results in the detection of different subsets of metabolites, precluded their side-by-side comparison (Table 1). For example, targeted mass spectrometry enables the precise determination of concentrations for a finite number of metabolites, whereas untargeted mass spectrometry detects up to two or three orders of magnitude more metabolites but is compositional in nature and thus does not yield absolute abundances. Furthermore, batch effects due to sample-processing such as differences in reagents, sample storage and mass-spectrometry instruments can introduce unwanted variation in both the abundances and the detected molecular features [26]. One additional obstacle we encountered was the proprietary nature of many of the metabolomic datasets that made it impossible to access the raw data and run standardized workflows.

Of the 40 transcriptomic datasets that were available in recount3 [100], the vast majority were obtained from studies with model animals, and only four of them had been obtained from postmortem processing of brain samples from autistic and neurotypical individuals. These four datasets collected different brain tissue types, including from the amygdala, the prefrontal cortex, the anterior cingulate and the dorsolateral prefrontal cortex.

### Data processing

16S amplicon and shotgun metagenomics samples were downloaded from the SRA. The 16S amplicon samples were processed using Deblur and subsequently mapped to bacterial whole genomes captured in the Web of Life using Woltka[101]. This is done in order to make the amplicon data comparable to shotgun metagenomics data. Bacterial abundances were extracted from shotgun metagenomics samples using Woltka and Bowtie2. Viral abundances were extracted from shotgun metagenomics samples using GPD and BWA. RNA expression data were obtained directly from recount3 [100]; the four metabolomics datasets were provided by the authors.

The approach of mapping both 16S amplicon sequences and SMS samples to a common set of microbial reference genome provided a consistent taxonomic annotatation between the different 16S amplicon types and SMS datasets. However, there are notable limitations in taxonomic resolution in all of these datasets. For instance, multiple Bifidobacterium species that are associated with non-human microbiomes were found including *B. asteroides* (honeybee), *B. callitrichos* (marmoset), *B. choerinum* (pig) and *B. sanguine* (tamarin). We observed that these 16S amplicons were multi-mapped across many different Bifidobacterium genomes, which highlights the lack of species level resolution highlighted in previous studies [102]. Similarly, taxonomic profiles obtained from shotgun sequencing are known to have elevated false positive rates taxonomic identifications due to high genome similarity between microbial reference genomes [59].

To enable age- and sexmatching, a bipartite matching between ASD and neurotypical subjects was performed using age and sex covariates. Subjects that could not be matched were excluded from the meta-analysis. Amongst the 16S and SMS datasets, there were multiple longitudinal datasets. To integrate these datasets into the cross-sectional analysis, we only picked the first time point for each subject.

### Differential ranking analysis

One of the most common approaches to evaluating microbiome and other omic studies consists of determining differences in the abundances of microbial taxa, human metabolites or other omic features between cases and controls [103]. Such differential abundance analysis is typically performed by computing the log fold changes between the case and control groups[104, 46, 105]. However, confounders such as sex-, age-, and geography-related batch effects, compositionality, high-dimensionality, over-dispersion, and sparsity, prevented a reliable estimation of differential abundances and thus compromised the side-by-side comparison of these differential abundances across studies in the manner of a traditional meta-analysis [106, 107]. Here, we set out to overcome these inherent limitations of traditional meta-analyses by developing a generalizable approach for controlling for select confounders that would help reveal a comprehensive picture of ASD-specific omic signals.

To minimize confounder effects, we developed a Bayesian differential ranking algorithm that used bipartite matching to optimize the age- and sex-based pairing of ASD and control subjects within each dataset. This approach helped both control for potential age and sex confounders and minimize batch effects such as sample collection method, sample processing protocol, and geographical provenance [108]. These Bayesian models were fitted via MCMC using Stan [109]. Conceptually, this allowed us to compute log-fold change differences of microbes between age- and sex-matched subjects, but because we did not have absolute abundance information we could only estimate this log-fold change up to a constant [31] (Supplemental methods). To determine if there was a significant difference between the age- and sex-matched pairs, we constructed an effect size metric utilizing our model’s uncertainty estimation (see Supplemental methods for more details). To show that this model is relevant for biological data, we built a simulation benchmark using the 16S count data. Specifically, we fitted the Bayesian model on the 16S cross-sectional cohort, and simulated microbial counts based on those estimated parameters. We showed that the ground truth log-fold changes across all of the microbes are within the 95% credible intervals estimated by our algorithm. When we evaluated our Bayesian model fit on the 16S, SMS and RNAseq datasets, our models fits achieved Rhat values below 1.1 and ESS values above 300, indicating that the draws from the posterior distribution are reliable [109].

To identify microbes that were ASD-specific or neurotypical specific, we fitted a Gaussian mixture model on top of the estimated log-fold changes, binning the taxa into three different groups, those taxa hypothesized to be more abundant in neurotypical controls, those more abundant in ASD children and those that are equally prevalent in both groups, or are “neutral”.

This strategy was inspired by the work done with ANCOM-BC [110]. The major difference in our approach compared to ANCOM-BC is that our approach assumes a negative binomial distribution for modeling counts and allows for Bayesian model uncertainty quantification. The reference frame in the cross-sectional analysis refers to the average abundance of the microbes that are categorized as neutral in Figure 1a. These same microbes were used to construct a reference frame in the FMT analysis to standardize all of the time points. The FMT analysis used the same matching strategy, but instead of matching on age and sex, the matchings were performed on the subjects to compare different time points.

The heatmap shown in Figure 1 displayed the log-fold changes for each case-control pair. To do this, a robust CLR transform was performed and all zeros were imputed to the mean abundance for visualization purposes. The case-control log-fold changes were computed for each pair as highlighted in Figure S7.

### Other methods

We fitted Random Forests models on nine 16S datasets and on three SMS datasets. We randomly split the samples into 90/10 training and test splits, performed a 10-fold cross-validation on the training datasets to obtain optimal model parameters, and computed predictions on the held-out test dataset. PERMANOVA with Bray-Curtis distances was used to determine if confounding variation due to household, age and sex were statistically significant in the sibling cohorts.

We used MMvec [111] to perform the diet-microbe co-occurrence analysis. Here, microbes were used to predict dietary intake. This analysis enabled the estimation of conditional probabilities, namely the probability of observing a dietary compound given the microbe was already observed. To estimate these conditional probabilities, MMvec performs a matrix factorization, identifying the factors that explain the most information in these interactions. We compared the MMvec microbial factors against the cross-sectional log-fold changes. We then compared the MMvec dietary factors against t-statistics that measure the differences in dietary compounds between ASD and neurotypical children.

To identify candidate viral-microbe interactions, we ran MMvec on each of the SMS datasets. We then pulled out the top co-occurring viral taxa for each microbe that had a conditional log-probability greater than 1, amounting to 78580 microbe-viral interactions. Then we filtered out the microbe-viral interactions that were not present in the GPD [70], leaving 31276 microbial-viral interactions.

We used Songbird [31] to perform the cytokine-microbe analysis via a multinomial regression that used the cytokines to predict microbial abundances. We reported biased microbial log-fold changes with respect to cytokine concentration differences. Pearson correlation was used to determine the agreement between the 16S cross-sectional microbial differentials and the microbe-cytokine differentials. To directly link these microbial abundances to the cytokine concentrations, we computed log-ratios, or balances, of microbes for each sample. For example, for IL-6 the numerator consisted of the top 30 microbes that are estimated to increase the most in abundance when IL-6 concentration increased, and the denominator consisted of the bottom 30 microbes which are estimated to be the most decreased when IL-6 concentration increases. Once these partitions are defined, the balances for each sample are computed by taking the log-ratio of the average abundance of the numerator group and the denominator group. See Morton et al 2017 for more details behind balances [112]. Pearson correlation between these balances and the cytokine concentrations are then computed to measure the agreement between the microbial abundances and the cytokine concentrations.

To identify key microbial genes, we performed a comparative genomic analysis in which we binned the microbial genomes into those associated with ASD and those associated with control subjects. Using a binomial test, we were able to determine if a particular gene was more commonly observed in ASD-associated microbes than by random chance. Significant microbial genes and RNA transcripts were subsequently mapped to KEGG pathways. To directly compare the two contrasting omics levels and gauge metabolic similarity, we retrieved all the molecules involved in both the microbial and human pathways and calculated their intersection. Since the metabolomics datasets were not directly comparable, we performed Wilcoxon tests on age- and sex-matched metabolomics samples within each cohort separately. While our analysis revealed multiple metabolites that were below the 0.05 p-value threshold, none of these metabolites passed the FDR corrected threshold.

## Acknowledgements

We would like to thank Allan Packer, Paul Wang, Natalia Volfovsky, Kelsey Martin and John Spiro for their critical review of the manuscript. We’d also like to add Kevin Liu, Hannah Sherman and Xue-Jun Kong for insightful discussions. Y.S. and T.D.L. are supported by the Wellcome Trust (WT098051).

## Software Availability

Software implementation of our Bayesian age-sex matched differential ranking algorithm can be found at https://github.com/flatironinstitute/q2-matchmaker

We want to acknowledge Matplotlib [113], Seaborn [114], Scipy [115], Numpy [116], Xarray [117], Arviz [118], Scikit-learn [119], biom-format [120] and Scikit-bio [121] for providing the software foundation that this work was built upon.

## Tables and Figures

**Figure BOX 1:**
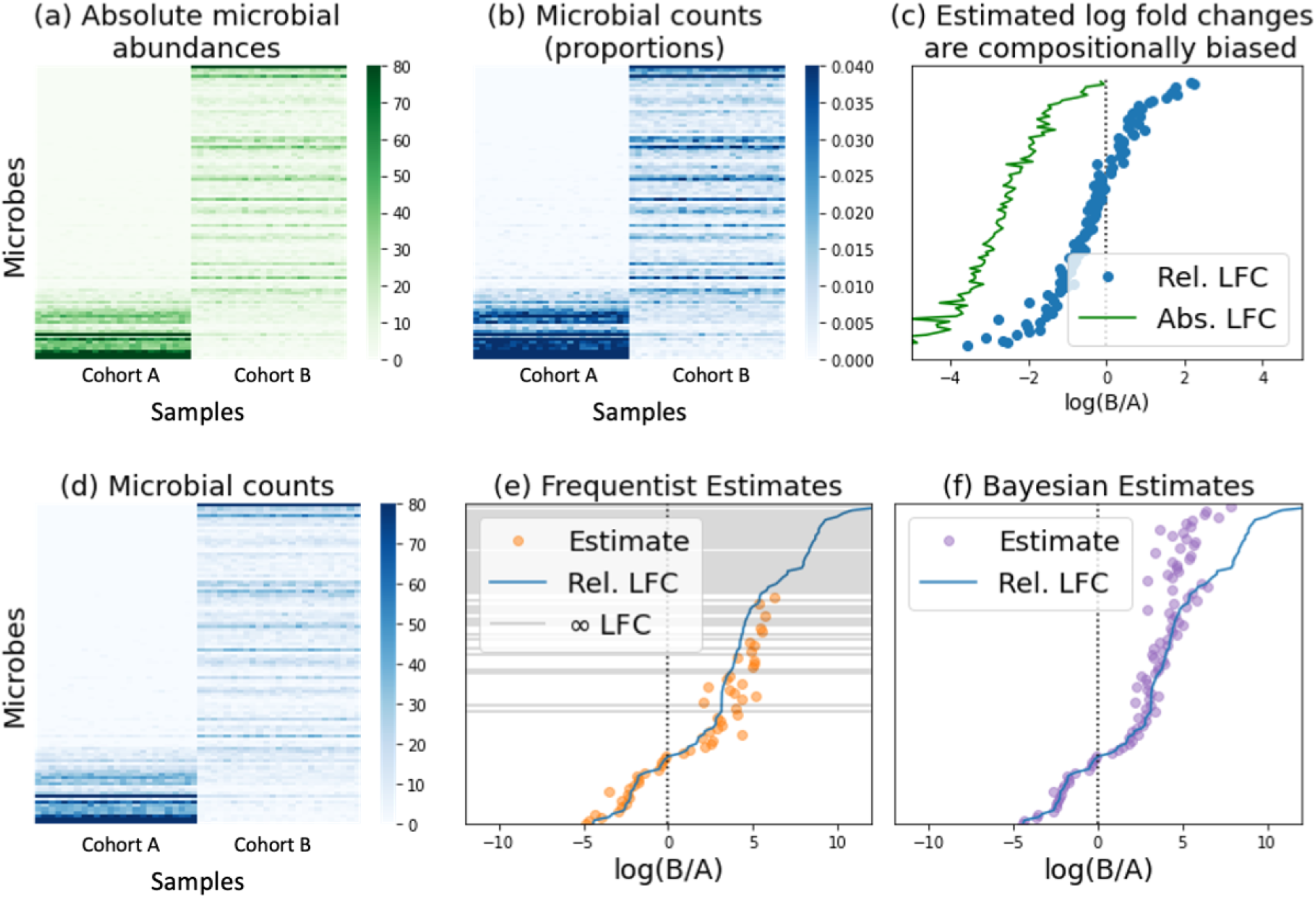
TACKLING METAGENOMIC UNKNOWNS. Metagenomic sequence data present unique quantification challenges due to a lack of total microbial load measurements, which precludes the determination of absolute microbe abundances, and to limitations brought about by sampling and sequencing depth limitations, which result in an incomplete representation of the metagenome. We devised a Bayesian differential ranking algorithm to address both these challenges, the compositional challenge and the zero-inflation challenge. **The compositional challenge**: Most sequencing count datasets lack absolute abundance information in the form of cells, colony forming units, or transcripts per volume. This limitation preempts the reliable estimation of log fold changes and is a defining characteristic of compositional data that can lead to excessive false positives or false negatives depending on the magnitude of the change in absolute abundances [31, 45]. As illustrated in panels a) through c), microbial counts (a) are typically converted into proportional abundances (b) that are then used to compute log-fold ratios. Fold change calculations adopt the general formula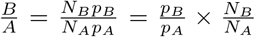 where *A* and *B* represent the two samples being compared, *pA* and *pB* represent the microbial proportions in *A* and *B*, and *N*_*A*_ and *N*_*B*_ represent the total number of microbes in *A* and *B*, also known as the ground truth. A key limitation of sequencing count data is their lack of proportionality to the corresponding absolute abundances in the original samples due to sequencing depth constraints [122]. Our inability to observe *N*_*A*_ and *N*_*B*_ introduces a bias that ultimately prevents us from performing FDR correction to identify differentially abundant microbes [123]. This bias depends on the change in microbial population size, with large population shifts leading to increased false positive and false negative rates, and an overall skewed representation of the ground truth (c). **The zero-inflation challenge**: Sampling errors and shallow sequencing lead to disproportionately high numbers of zero counts, especially for microbes present in low abundances (d). Multinomial, Poisson and Negative Binomial distributions have been used to explicitly handle zero counts [46]. However, estimating log-fold differentials remains problematic when microbes are not observed in any of the samples in one group since log 0 is −∞ and thus the true log-fold change of a zero-count microbe can not be determined (e). Bayesian inference avoids this problem by introducing a prior that prevents nonsensical log-fold change estimates (f). Specifically, this introduces a rounded-zero assumption whereby all microbes have a non-zero chance of being observed. Panel h highlights what these log-fold changes would look like using a Dirichlet prior, where every microbe has the same probability of being observed before collecting data.

## Supplemental Materials

**Figure S1:**
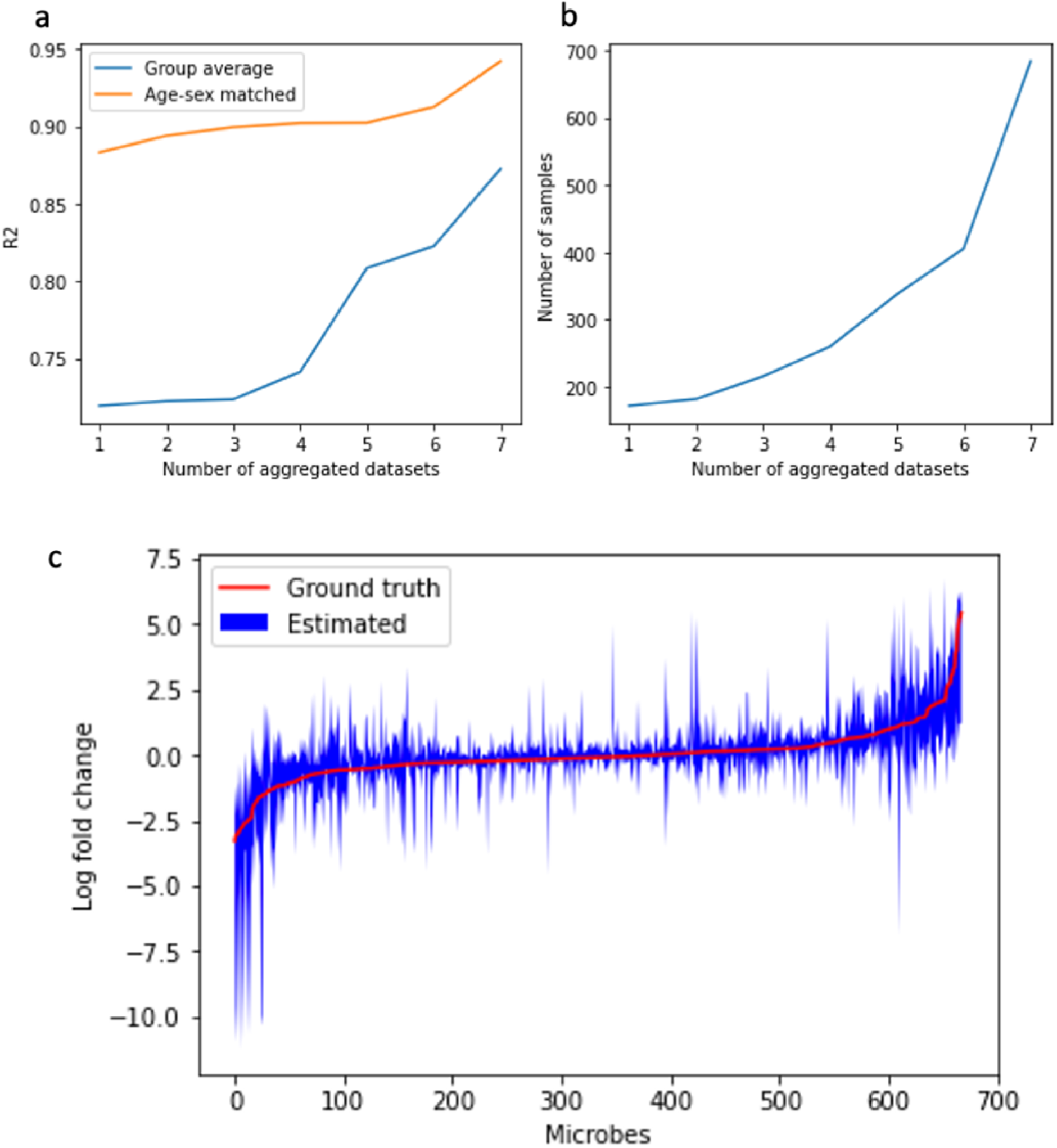
Benchmarks. (a) Comparison of age- and sex-matching approach compared to standard group averaging with respect to dataset size across 7 of the 10 16S studies (excluding Kang et al, David et al and Son et al). (b) Number of samples analyzed. The x-axis represents the number of aggregated datasets, the y-axis on the left panel is the average *R*^2^ metric to measure the model error, and the y-axis on the right panel is the number of samples in the aggregated dataset. (c) Differential abundance estimation derived from a simulated datasets modeled from the cross-sectional cohort from the 8 16S datasets.

**Figure S2:**
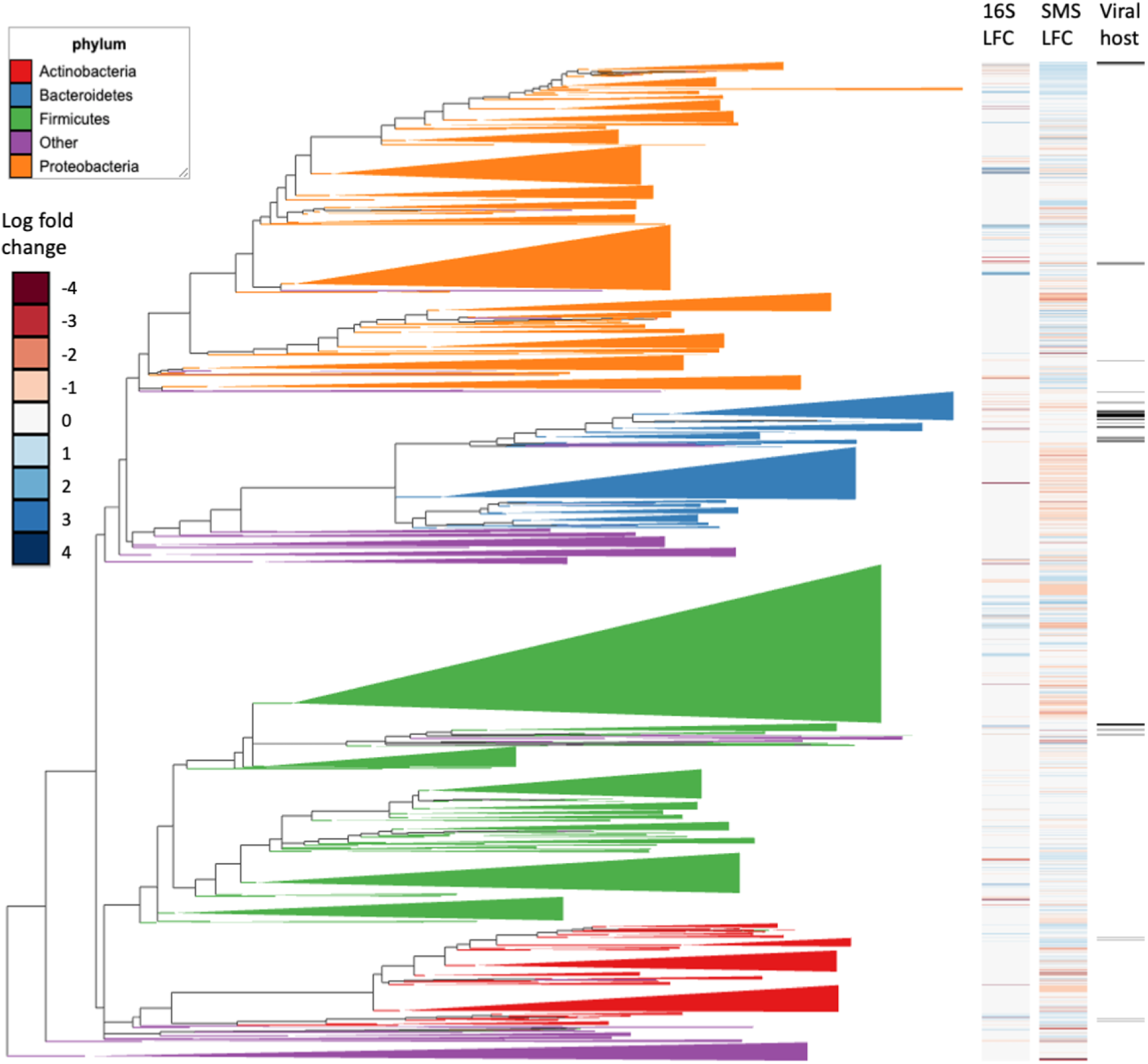
Phylogenetic visualization of microbes with respect to the differentials computed from 16S and SMS differentials. Microbes that were annotated as viral hosts were also highlighted. The phylogenetic visualization was generated using Empress [125].

**Figure S3:**
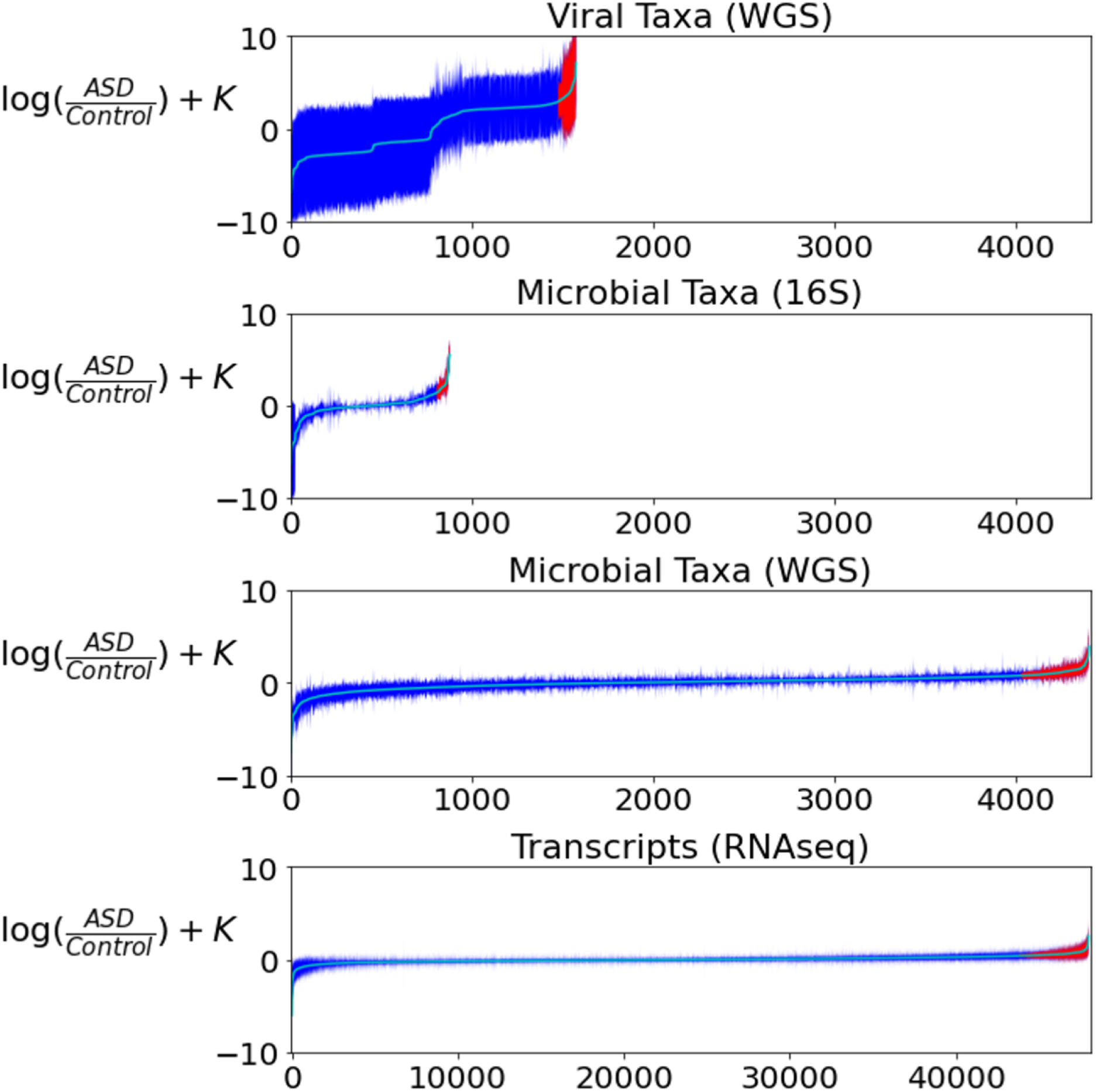
Global differential ranking trends observed for the virome, 16S, SMS, and RNAseq datasets analyzed in this study. The x axis for the virome, 16S and SMS datasets is equivalent to showcase the differences in feature counts; the x axes for the RNAseq dataset is larger by a factor of 10, illustrating the stark difference in number of features of this dataset compared to the other three.

**Figure S4:**
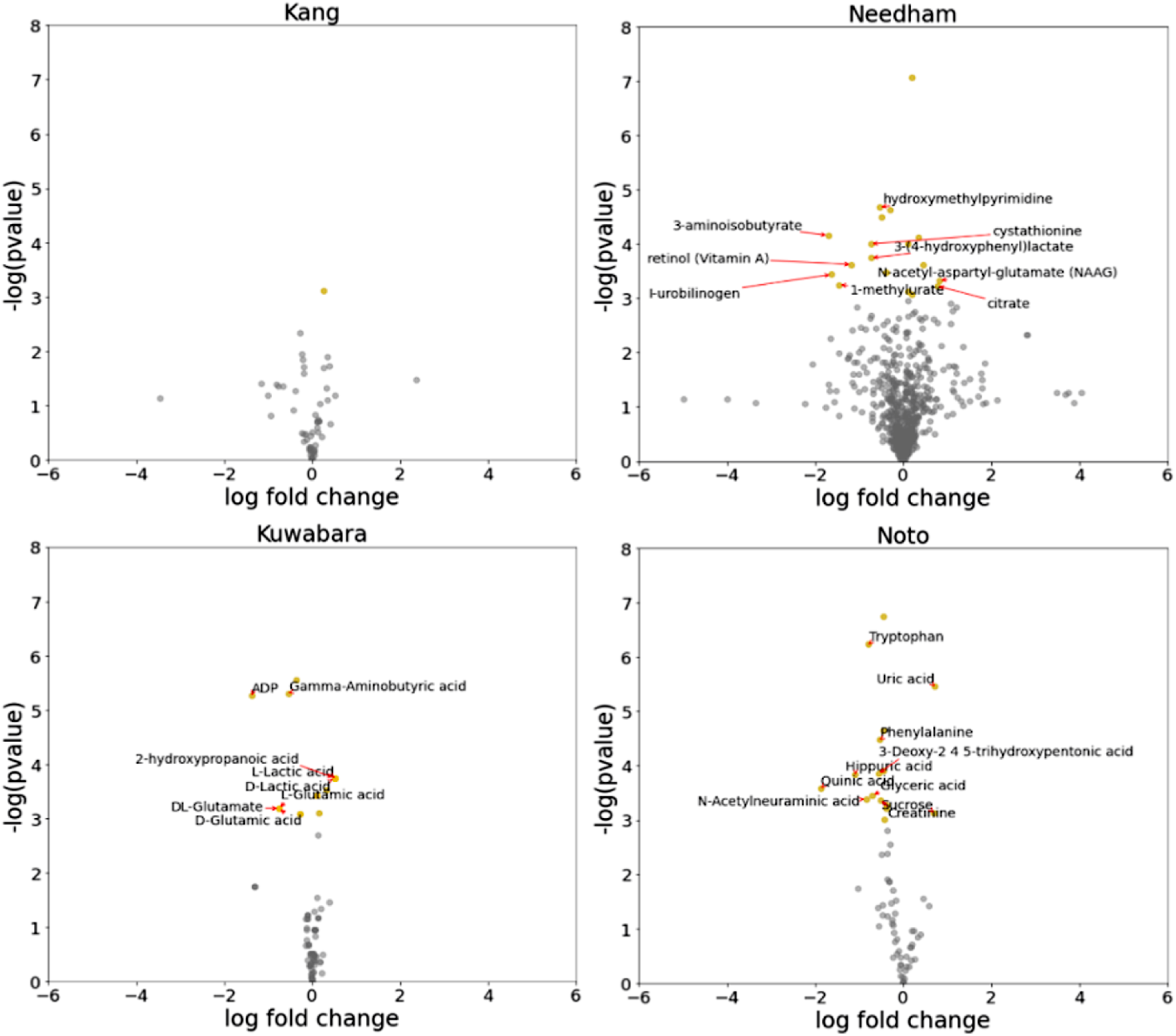
Metabolomics differential ranking analysis across four studies. Paired t-tests were performed to identify differentially abundant metabolites. The metabolites shown in Needham et al consist of both fecal and serum metabolites. None of the metabolites had significant log-fold changes after applying FDR correction.

**Figure S5:**
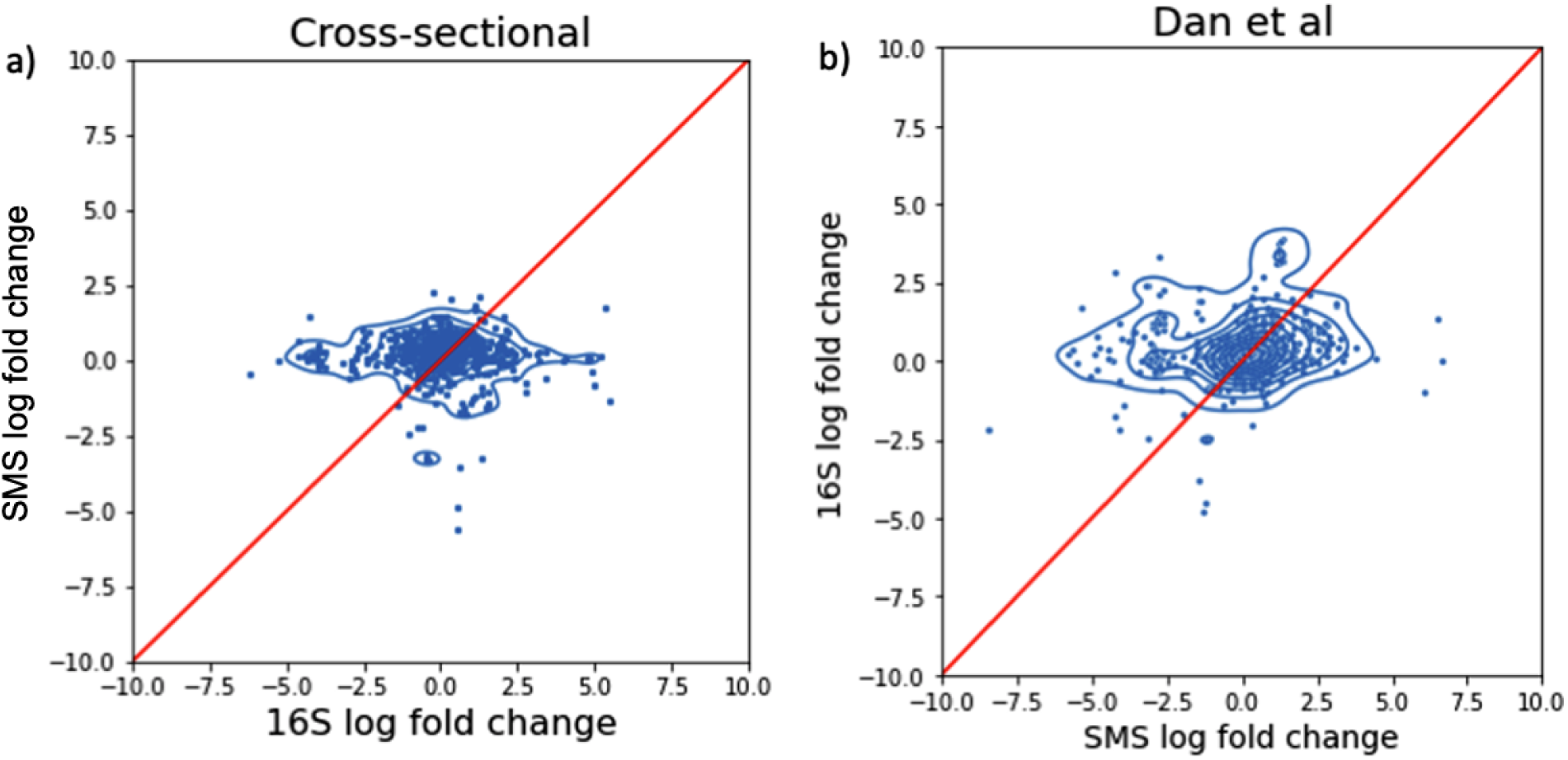
Comparison of log-fold changes computed from 16S and SMS. (a) Comparison of differentials obtained from 16S and SMS on the cross-sectional datasets. (b) Comparison of differentials obtained from 16S and SMS on the same samples from Dan et al.

**Figure S6:**
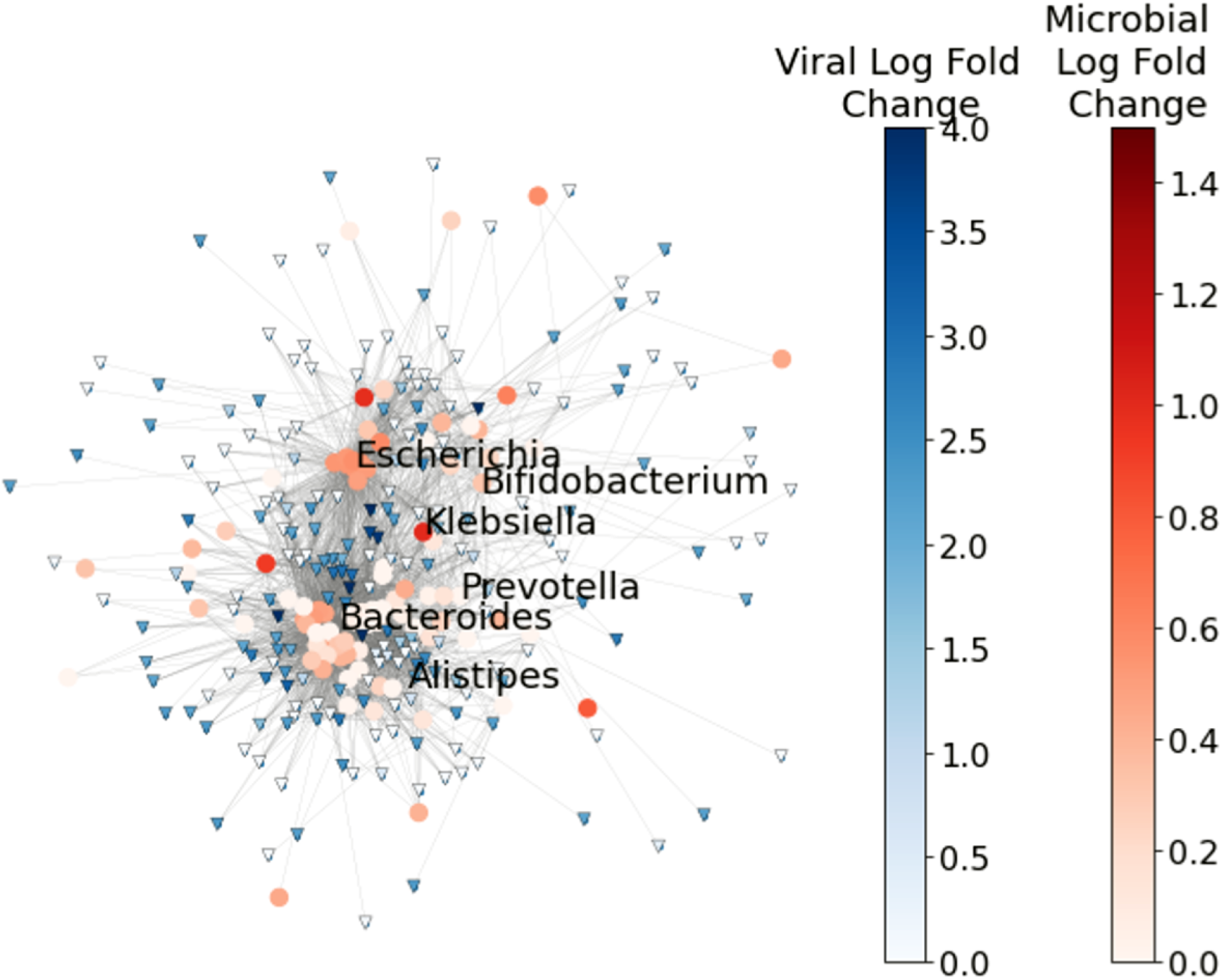
Microbe-viral co-occurrence network estimated using MMvec. Microbes are colored red and viruses are colored blue. Edges are drawn between microbes and viruses if they are highly co-occurring and the interaction was annotated in GPD.

**Figure S7:**
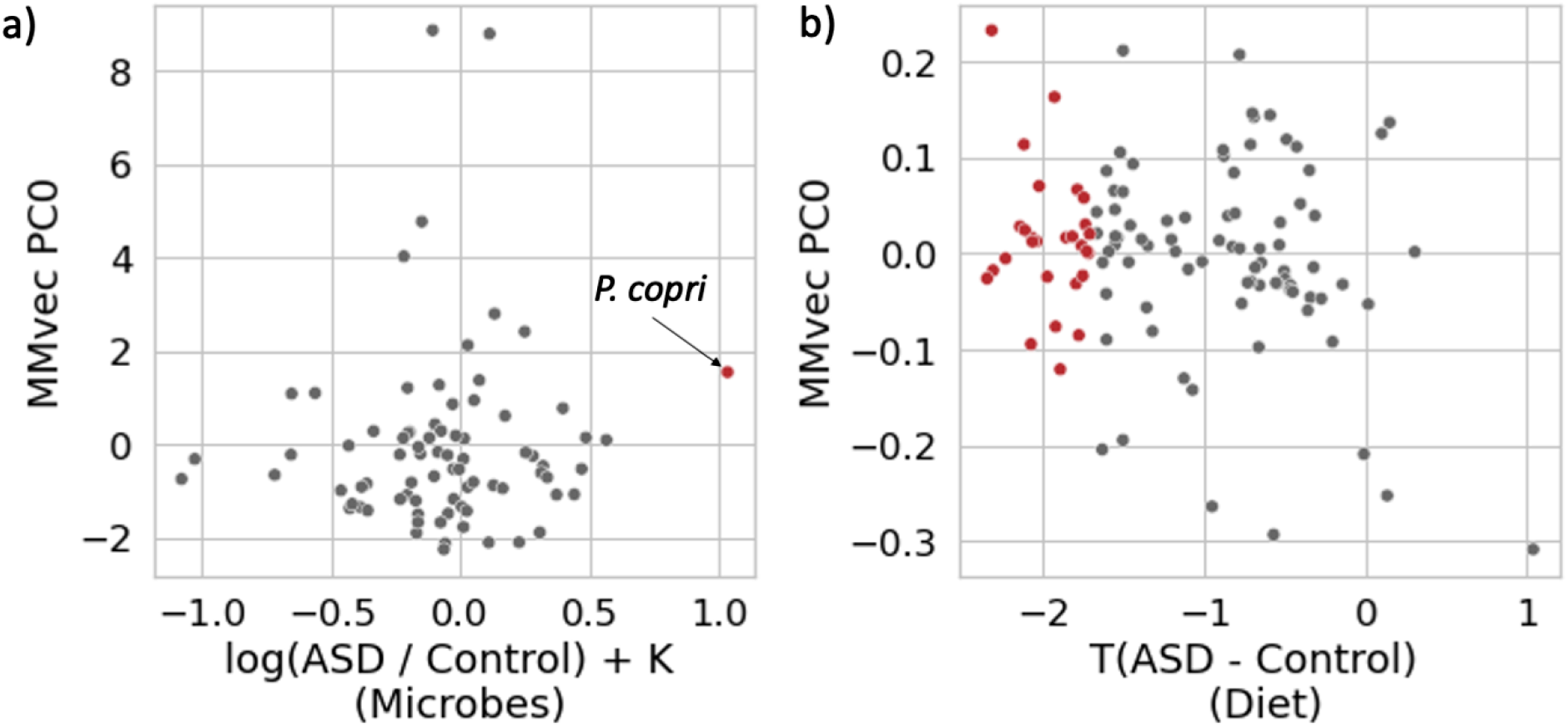
Principal components analysis of microbe-diet interactions. The top principal component explaining the variation in the microbe-diet co-occurrences is compared against the (a) microbial log fold change and (b) dietary differences computed from a t-statistic. Dietary compounds that are significantly significant before FDR correction are highlighted in red.

**Figure S8:**
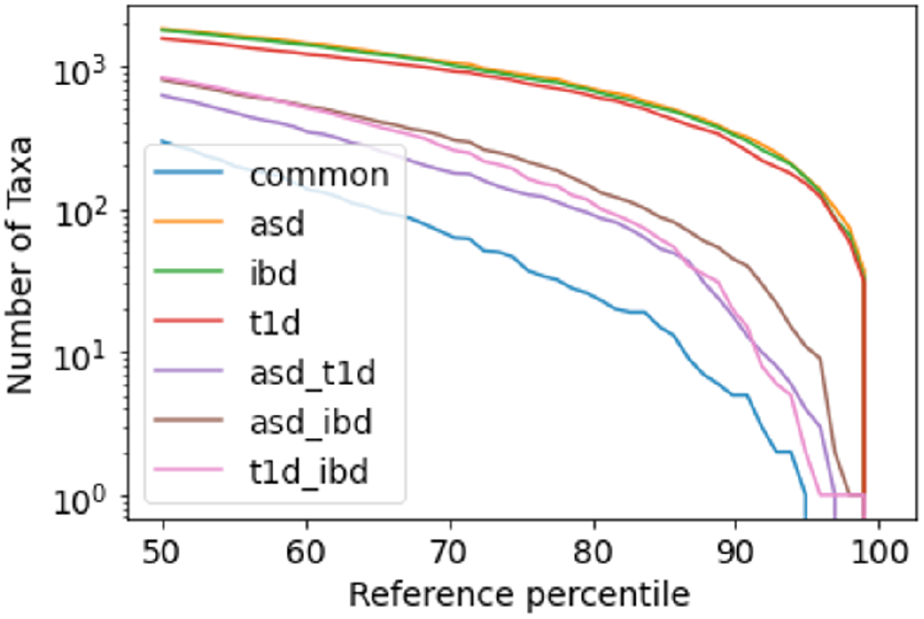
Comorbidity analysis. The number of taxa in common between diseases from differential ranking analysis are shown, with a focus on the intersections between ASD, IBD and T1D.

Table S1: Table of statistics for 16S differentials, including mean log-fold change, standard deviation log fold change, 95% credible intervals and taxonomy for each microbe.

Table S2: Table of statistics for SMS differentials, including mean log-fold change, standard deviation log fold change, 95% credible intervals and taxonomy for each microbe.

Table S3: Table of statistics for RNAseq differentials, including mean log-fold change, standard deviation log fold change, 95% credible intervals and taxonomy for each transcript.

Table S4: Table of statistics for viral differentials, including mean log-fold change, standard deviation log fold change, 95% credible intervals for each virus.

Table S5: Permanova breakdown of sibling matched cohorts looking at the confounding variation due to age, sex and household.

Table S6: Microbial log fold changes due to cytokine differences, including mean log-fold change for each cytokine.

Table S7: Microbe virus co-occurrences, where entries represent the centered log-probability of a microbe and a virus both present for a given sample

Table S8: A list of paired microbe and human pathways in addition to the number of overlapping metabolites

Table S9: Microbe diet co-occurrences, where entries represent the centered log-probability of a microbe and a dietary compound both present for a given subject.

Table S10: Microbial log fold changes between paired time points across all of the subjects in the FMT study.

## Notes

**Conflict of Interest** R.H.M. is Scientific Director at Precidiag Inc.; T.D.L. is co-founder and Chief Scientific Officer of Microbiotica; S.K.M. is a co-founder and has equity in Axial Therapeutics; R.B is currently Executive Director of Prescient Design, a Genentech Accelerator; G.T.-O. is a Consultant-in-Residence at the Simons Foundation.

### Competing Interest Statement

R.H.M. is Scientific Director at Precidiag Inc.; T.D.L. is co-founder and Chief Scientific Officer of Microbiotica; S.K.M. is a co-founder and has equity in Axial Therapeutics; R.B is currently Executive Director of Prescient Design, a Genentech Accelerator; G.T.-O. is a Consultant-in-Residence at the Simons Foundation.

