## Supplemental Materials for "Multi-omic analysis along the gut-brain axis points to a functional architecture of autism"

February 26, 2022

### 1 Bayesian Differential Ranking

Conceptually, the goal of a differential analysis is to make a statement about change in abundance for a given feature $i$ between conditions A and B by evaluating the following null hypothesis:

$$\frac{A_i}{B_i} = 1$$

However, most omic datasets do not provide a direct observation of the absolute quantities of $A_i$ and $B_i$, or the total microbial loads $N_{A_i}$ and $N_{B_i}$, but rather only an observation of their proportions $p_{A_i}$ and $p_{B_i}$, respectively, within each dataset, which are determined by a bias term, $\frac{N_A}{N_B}$. This bias term, given by

$$\frac{A_i}{B_i} = \frac{p_{A_i} N_A}{p_{B_i} N_B} = \frac{p_{A_i}}{p_{B_i}} \frac{N_A}{N_B}$$

results in high false discovery rates (FDRs) that cannot be adjusted for in models analyzing compositional omics datasets because the overall contribution of $N_A$ and $N_B$ to change can not be unequivocally quantified (Vandeputte et al. 2017; Hawinkel et al. 2019). To avoid the total biomass bias without having to resort to performing traditional FDR corrections, we adopted a ranking approach that allowed us to sort omic features by their log-fold change values independently of how large their change was in absolute terms (Morton, Marotz, et al. 2019). Since the biomass bias impacts every species within a dataset equally, the ranking approach ignores this bias, making the approach scale-invariant (Equation 1).

$$\text{rank}(\frac{A}{B}) = \text{rank}(\frac{p_A N_A}{p_B N_B}) = \text{rank}(\frac{p_A}{p_B}) \tag{1}$$

The overall model we designed consisted of a customized differential abundance tool that leveraged the experimental design of each study included in the analysis to determine study-specific feature perturbation profiles that could then be combined with the normalized perturbation profiles of other studies to perform a global differential perturbation analysis. The overall model had the following structure,

$$y_{i,j} \sim \text{NegativeBinomial}(\lambda_{i,j}, \alpha_{ij})$$
$$\log \lambda_{i,j} = \log N_i + C_{k(i),j} + D_j \text{I}[i = ASD]$$

where $y_{i,j}$ denotes the microbial counts in sample $i$ of species $j$ across $d$ species. $\lambda_{i,j}, \alpha_{ij}$ represents the expected counts for species $j$ and sample $i$, $j$ represents a microbe specific over-dispersion term, $N_i$ represents sequencing depth (self normalization and preemptive of rarefaction), $C_{k(i),j}$ represents the log proportion of species $j$ in the $k(i)$ control subject (age- and sex-matched), and $D_j \text{I}[i = ASD]$ represents the log-fold change difference between control and ASD subject with a corrective function that equals 1 when $i$ corresponds to the paired ASD subject and 0 when $i$ corresponds to the control subject. Incorporating $N_i$ into the model renders the model self normalizing and not dependent on rarefaction, and $C_{k(i),j}$ incorporates the age- and sex-matching component for a given pair $k$.

The priors for these variables are given below

$$\alpha_{ij} = \frac{a_{0,k(i),j}}{\lambda_{ij}} + a_{1,k(i),j} + \beta_p \qquad a_0 \sim \text{LogNormal}(0,1) \qquad a_1 \sim \text{LogNormal}(\log(10), 0.1)$$
$$\beta_p \sim \text{Normal}(\beta_\mu, \beta_\sigma) \qquad \beta_\mu \sim \text{Normal}(0,3) \qquad \beta_\sigma \sim \text{LogNormal}(\log(0.1), 0.1)$$
$$C_{k(i),j} \sim \text{Normal}(C_{\mu_j}, C_{\sigma_j}) \qquad C_{\mu_j} \sim \text{Normal}\left(\frac{1}{d}, 3\right) \qquad C_{\sigma_j} \sim \text{Normal}\left(\frac{1}{d}, 1\right)$$
$$D_j \sim \text{Normal}(0,3)$$

Here, the overdispersion parameter are estimated for each microbe, each batch as well as the ASD and control groups. This approach is adapted from DESeq2, allowing for the overdispersion to be modeled by both linear and quadratic terms with respect to the abundance. Furthermore, this parameterization does allow for a compositional interpretation due the following rationale; The Poisson distribution with an offset term is known to approximate the Multinomial distribution [1, 2, 3, 4]. Furthermore, the Negative Binomial can be reparameterized as a Gamma-Poisson distribution, allowing for overdispersion modeling by breaking the mean, variance relationship inherient in the Poisson distribution.

The age-sex matched differential abundance has a similar methodology to paired tests such as paired t-test and Wilcoxon test. To this end, we also used this differential abundance methodology to analyze the FMT dataset. Here instead of matching pairs of subjects, we matched pairs of time points and computed the differential abundance across each pair of time points. To make these differentials comparable, a common set of taxa that were detected to be associated with controls were selected. Specifically, taxa that had a log-fold change less than 0 in the cross-sectional cohort were assigned to this reference set. The estimated log-fold changes are adjusted by centering around the mean log-fold in the reference dataset as follows

$$D^* = D - \bar{D}_R$$

where $\bar{D}_R$ denotes the mean of the log-fold changes of the reference taxa, and $D^*$ represented the recentered log-fold changes. By doing this, all of time points will have the same reference and will be more directly comparable.

One of the advantages of the above model is that it will cancel out any multiplicative batch effect such as PCR amplification bias with no impact on j. This is because $D$ is only computed within cohorts and as a result, cohort specific batch effects are mitigated. Furthermore, this differential abundance model can be applied to different types of omics data. And because we built the differential ranking model in a Bayesian environment, we were able to fit the model using an MCMC approach to estimate uncertainty by sampling the resulting posterior distributions.

For example, to make a statement about the value of an estimated posterior probability distribution $p(D|y)$ we could compute an average value using the following approximation:

$$\mathbb{E}[D] \approx \frac{1}{m}\sum_{i=0}^{m} \hat{D}_i \qquad \hat{D}_i \sim p(D|y)$$

Using this classic application of MCMC sampling in which N samples of i are drawn from the posterior distribution $p(D|y)$, we were able to approximate the true mean of the posterior differential abundance distributions and the corresponding effect sizes. With this, we can compute an effect size metric that determines if there is any global difference detected. This metric is analogous to PERMANOVA, but one that computes this from log-fold changes using the age-sex matched design. The effect size $E$ is measured as follows:

$$E = \frac{||\mu_D||_2}{r_D}, \qquad \mu_D = \frac{1}{m}\sum_{i=0}^{m} \hat{D}_i \qquad r_D = \max_{\hat{D}_i \sim p(D|y)} |||\hat{D}_i - \mu_D|||_2 \tag{2}$$

where $\mu_D$ is the mean of the posterior distribution and $r_D$ represents the radius of a sphere that contains all of the samples from the posterior distribution. If the effect size is greater than 1, that means that zero is not included in the posterior distribution and the difference is significant. Bayesian p-values are computed from the number of draws of $\hat{D}_i$ that were simulated from the posterior distribution $p(D|y)$. For instance if 100 draws are sampled from the posterior distribution, and zero is not within the sphere estimated from those 100 draws, then we say that the posterior distribution is significantly not overlapping with zero with pvalue $< 0.01$.
